# Phytohormone profiling in an evolutionary framework

**DOI:** 10.1101/2023.04.06.534998

**Authors:** Vojtěch Schmidt, Roman Skokan, Katarina Kurtović, Stanislav Vosolsobě, Roberta Filepová, Samuel Haluška, Petre Dobrev, Václav Motyka, Jan Petrášek

## Abstract

Multiple phytohormones act as conserved developmental regulators in land plants. Although the closely related streptophyte green algae typically lack full complements of molecular pathways underlying these responses, scattered reports of endogenous phytohormone production in these organisms exist. In this study, we performed a detailed LC/MS-based analysis of several phytohormones, their precursors and metabolites in all lineages of streptophyte algae. We also included chlorophyte algae and early-diverging land plants as outgroups. Free auxin, tRNA-derived cytokinins and certain phenolics including salicylic acid were found ubiquitously. However, land plants differed from green algae by the consistent detection of abscisic acid and the presence of auxin and cytokinin conjugates and *trans*-zeatin, supporting the hypotheses that these three phytohormones likely came to regulate development in the ancestral land plant. By contrast, we observed a patchy distribution of jasmonates among streptophytes. We additionaly analyzed the corresponding culture and empty media to account for phytohormone excretion and environmental contamination. Extracellular auxins and cytokinins were frequently detected, while agar constituted a major external source of phenolic compounds. We provide a highly comprehensive evolution-directed screen of phytohormone compound occurrence and thoroughly discuss our data in the context of current plant hormonomics and phylogenomics.

**GRAPHICAL ABSTRACT:** 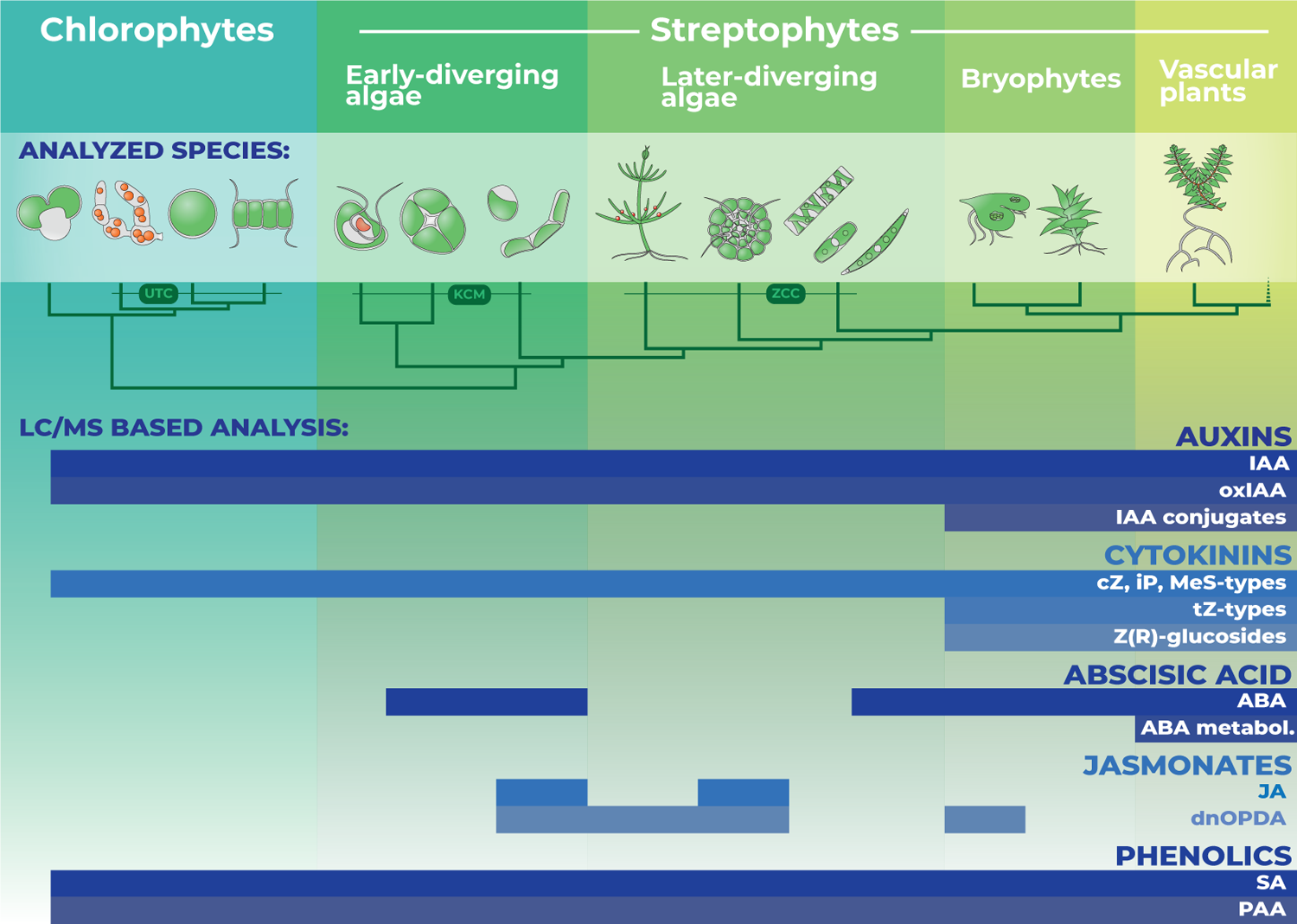

## INTRODUCTION

Land plants evolved from within a group of streptophyte algae, also called charophytes (Bowman, 2022). The extant charophytes are highly morphologically divergent, both mutually and from land plants (de Vries & Rensing, 2020), and range from unicellular to filamentous and even multicellular bodies harboring different cell types (Delwiche, 2016). The genomes of several representative charophyte species have recently been sequenced (Hori et al., 2014; Nishiyama et al., 2018; Cheng et al., 2019; Jiao et al., 2020; Liang et al., 2020; Wang et al., 2020; Sekimoto et al., 2023) and transcriptome databases of many more have become available (1kp, 2019; Cheng et al., 2018). This development has revealed that the molecular background underlying many characteristic traits of land plants had at least partially originated within charophytes, including response to several phytohormones (Rensing, 2018 and 2020; Furst-Jansen et al., 2020).

Phytohormones comprise several types of endogenous, low molecular weight compounds that elicit physiological and growth responses in cells and tissues where they are accumulated (Davies, 2010; Waadt, 2020; Sageman-Furnas & Strader, 2022; Waadt et al., 2022). Studies in bryophytes, the monophyletic sister lineage to vascular plants (Puttick et al., 2018; Harris et al., 2020; Su et al., 2021), revealed that many molecular pathways underlying phytohormone responses are functionally conserved between bryophytes and angiosperms (Guillory & Bonhomme, 2021), and thus were likely present in the common ancestor of all extant land plants. These include auxin biosynthesis, parts of metabolism, signaling and transport (Viaene et al., 2014; Bennett et al., 2014; Eklund et al., 2015; Flores-Sandoval et al., 2015; Kato et al., 2017 and 2018; Mutte et al., 2018; Thelander et a., 2018; Suzuki et al., 2022), cytokinin biosynthesis and signaling (Yevdakova & von Schwartzenberg, 2007; Coudert et al., 2019; Rashotte, 2021), abscisic acid biosynthesis and signaling (Takezawa et al., 2015; Sun et al., 2020), strigolactone biosynthesis and strigolactone/karrikin signaling (Proust et al., 2011; Lopez-Obando, 2021; Kodama et al., 2022) and the signaling of jasmonates (Monte et al., 2018; Penuelas et al., 2019), salicylic acid (Jia et al., 2023) and ethylene (Yasumura et al., 2012 and 2015). By contrast, gibberellin and brassinosteroid signaling emerged in vascular plants and euphyllophytes, respectively (Wang & Mao, 2014; Hernandez-Garcia et al., 2021; Zheng et al., 2022). In charophyte genomes, orthologs of these pathways are often absent or only partially and/or inconsistently present among taxa. To date, ethylene signaling is the only demonstrated example of a functionally conserved phytohormone response between land plants and charophyte algae (Ju et al., 2015). Nevertheless, endogenous phytohormones and various physiological responses to applied phytohormones have been recorded in green and other algae, particularly unicellular chlorophytes relevant to bioindustry (Han et al., 2018; Stirk & van Staden, 2020). However, these are only distantly related to land plants (Turmel & Lemieux, 2018; Bowles et al., 2022), with their close charophyte algal relatives receiving comparatively much less attention.

In this study we complement the recent genomic exploration of charophytes with a comprehensive profile of the occurrence of multiple phytohormones and their metabolites, additionally including several chlorophyte algae and early-diverging land plants as outgroups. We compare our data to the previously published evidence and discuss these in relation to the current state of phylogenomics.

## RESULTS AND DISCUSSION

### Experimental setup

The algal and land plant strains were selected based on their establishment in research, axenicity and ease of cultivation (see **Table S1** for details). The phytohormone measurements in biomass were supplemented with the same analysis in the corresponding culture media, to address possible excretion into the environment. In an, to our knowledge, unprecedented step we related these data to the same analysis performed in blank media without biological material to account for possible environmental contaminants (**Box 1**). We employed a generalized LC/MS-based approach designed to detect the broadest possible spectrum of phytohormones, but particularly the active forms, precursors and metabolites of auxins, cytokinins, abscisic acid, jasmonates and phenolics including salicylic acid. See **Table S2** for the full list of analytes and **Table S3** for the entire raw dataset.

**Box 1.**
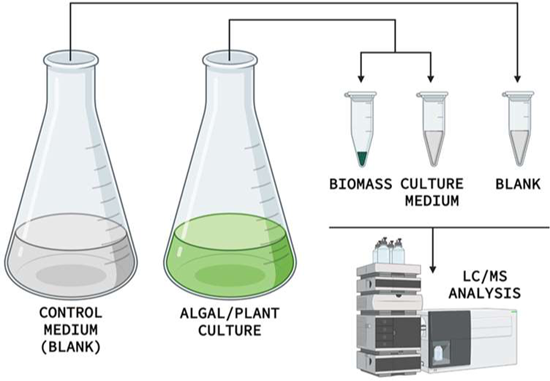
Experimental setup. Samples of biomass and the corresponding culture medium were taken. Additionally, samples of blank medium (containing no biological material and kept in the growth chamber alongside the growing culture) were taken to account for environmental contaminants. *(Created with BioRender.com)*

### Phytohormone levels in biomass

The profile of phytohormones and their metabolites measured in algal and land plant biomass is summarized in **Figure 1** (detailed version listing individual compounds in **Figure S1**) and discussed below.

**Figure 1.**
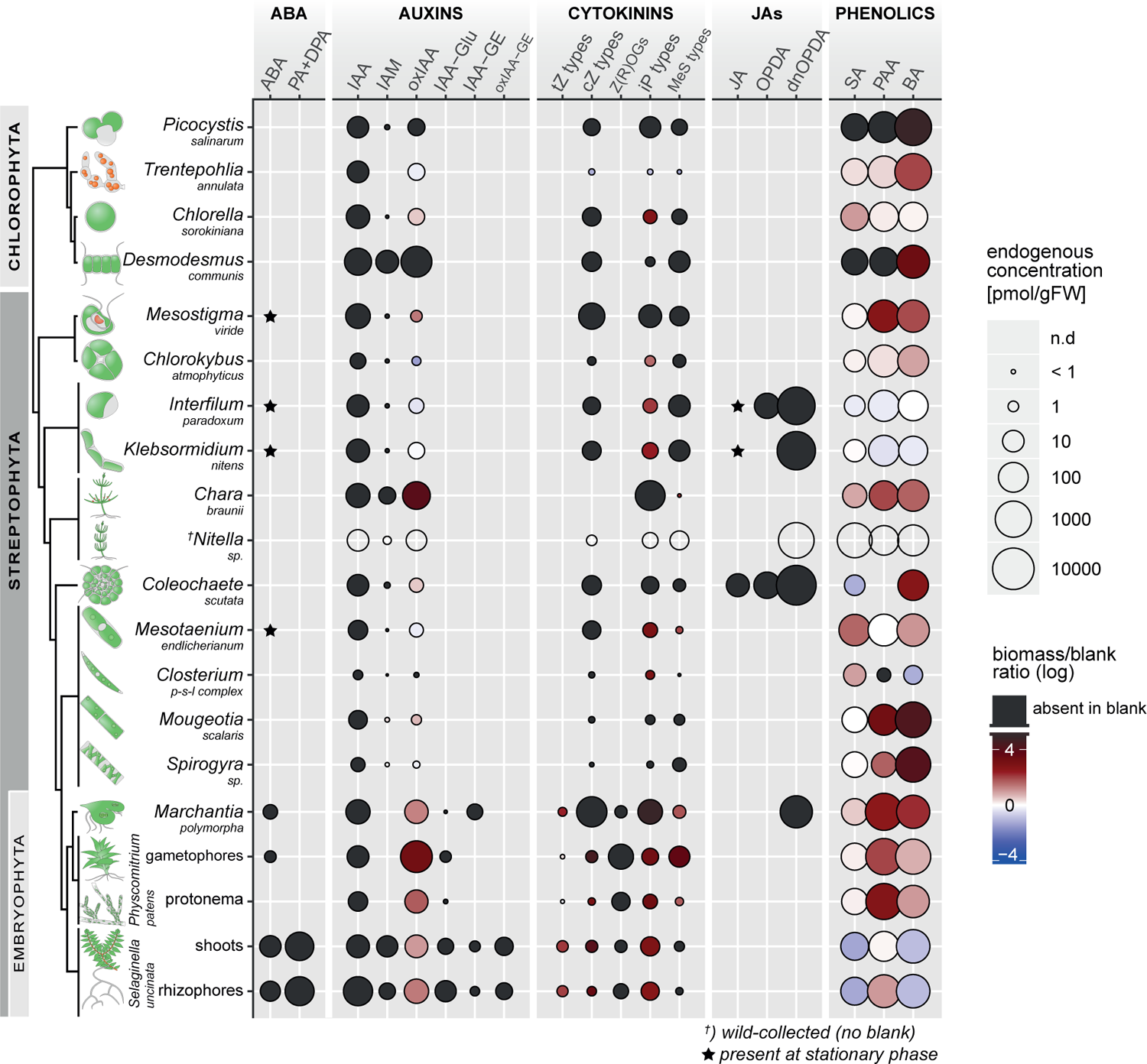
Endogenous phytohormone compounds detected in the biomass of green algae and land plants. Circle size denotes concentration in biomass (pmol per gram fresh weight). No circle–compound not detected (n.d.). Color code denotes the ratio between the concentrations measured in biomass and blank medium (the latter containing no biological material), expressed in logarithmic scale. Blue–compound(s) prevalent in blank. Red–compound(s) prevalent in biomass. Black–compound(s) absent in blank. Symbols: Star–compound only detected in stationary-phase cultures. Cross–wild-collected biological material (no blank available). Abbreviations: ABA (abscisic acid), PA (phaseic acid), DPA (dihydrophaseic acid), IAA (indole-3-acetic acid), IAM (indole-3-acetamide), oxIAA (2-oxo-IAA), IAA-Glu (IAA-glutamate), IAA-GE (IAA-glucose ester), tZ (*trans*-zeatin), cZ (*cis*-zeatin), Z(R)OGs (zeatin (riboside)-O-glucosides; both *cis-* and *trans-* isomers), iP (N6-(Δ 2-isopentenyl)-adenine), MeS (methylthio), JA (jasmonic acid), OPDA (12-oxo-phytodienoic acid), dnOPDA (dinor-OPDA), SA (salicylic acid), PAA (phenylacetic acid), BA (benzoic acid).

Abscisic acid (ABA) presence has been reported in chlorophyte and charophyte green algae (Hirsch et al., 1989; Tietz et al., 1986 and 1989; Marsalek et al., 1992; Hackenberg & Pandey, 2014; Hori et al., 2014; Stirk et al., 2014; Beilby et al., 2015; Chokshi et al., 2017) and other organisms (Hartung et al., 2010; Lu et al., 2014; Noble et al., 2014). By contrast, we convincingly detected ABA in only a few charophyte strains, in all cases stationary (older) cultures (**Figure S2A**), but not in any chlorophytes. An increase in endogenous ABA was previously observed in stationary cultures of the chlorophyte *Chlorella vulgaris* compared to proliferative ones (Marsalek et al., 1992), whereas we detected no ABA in *Chlorella sorokiniana* at either growth stage. By contrast, we did find ABA in all land plant samples except *Physcomitrium patens* protonema, which is a contentious issue in literature (Xiao et al., 2018; Rathnayake et al., 2018). Only *Selaginella* contained the ABA catabolites phaseic and dihydrophaseic acids (PA & DPA) in our dataset. However, these were previously found in wild-collected bryophytes (Zaveska Drabkova et al., 2015) and could be produced by the moss *Riccia* when supplied with exogenous ABA (Hellwege & Hartung, 1997). The concentrations of ABA we measured are largely comparable (within the order of magnitude) to those reported in bryophytes and *Selaginella* (Hellwege & Hartung, 1997; Liu et al., 2008; Wang et al., 2010; Kageyama et al., 2015; Xiao et al., 2018; Pizarro et al., 2019) or lower (Rathnayake et al., 2018; Zaveska Drabkova et al., 2015).

Auxin profile in our dataset was characterized by the omnipresence of free indole-3-acetic acid (IAA), its oxidized form (2-oxo-IAA; oxIAA) and indole-3-acetamide (IAM). Land plants additionally possessed amide- and ester-conjugates of IAA, namely IAA-glutamate (IAA-Glu) and IAA-glucose (IAA-GE). This mostly fits the available literature. oxIAA and IAM are frequently reported together with IAA in green algae (Jiraskova et al., 2009; Stirk et al., 2013 and 2014; Zizkova et al., 2017) and land plants alike (Pollmann et al., 2002; Novak et al., 2012; Vankova et al., 2017; Kaneko et al., 2020; Zlobin et al., 2020; Cosic et al., 2022; Fuchs et al., 2022; Prerostova et al., 2021). oxIAA has been studied mainly as an auxin catabolite in angiosperms where it is likely not produced directly from IAA (Hayashi et al., 2021); the upstream metabolites are not found in algae (including this study) and the full complement of known enzymes leading to oxIAA production seems to be angiosperm-specific (Bowman et al., 2021). Although IAA can oxidize into oxIAA under certain *ex vivo* conditions (Hu & Dryhurst, 1997; Kai et al., 2007), it is unlikely to be an artifact as the control samples containing pure IAA standards showed <1% contamination with oxIAA (data not shown), which is far below the ratio measured in biomass. Hence, endogenous non-decarboxylative IAA oxidation must be widespread, regardless of the mechanism. IAM is a known IAA precursor in plant-associated, typically pathogenic (Manulis et al., 1994; Costacurta & Vanderleyden, 1995; Maor et al., 2004; Cerboneschi et al., 2016) but also symbiotic (Tsavkelova et al., 2007; Gutierrez et al., 2009; Tsavlekova et al., 2012) bacteria and fungi, and marginally also in Arabidopsis (Gao et al., 2020). Although a land plant-specific two-step pathway accounts for the majority of auxin biosynthesis in land plants (Zhao et al., 2012; Eklund et al., 2015), an alternative represented by the AMIDASE1 (AMI1) enzyme (converts IAM to IAA) is conserved across the green lineage (Mano et al., 2010; de Smet et al., 2011; Liang et al., 2020), although this has not been studied further. Among auxin conjugates, IAA-Asp is often being included in profiling and usually is not detected in green algae (Stirk et al., 2013; Hackenberg & Pandey, 2014) and sometimes not even in bryophytes (Kaneko et al., 2020). When found, IAA-Asp is present in trace amounts, typically in only a few of several strains examined (Stirk et al., 2014; Zizkova et al., 2017). Such concentrations unfortunately would not be detectable in our analysis, because complications with an internal standard reduced the sensitivity of IAA-Asp detection. Finally, the phenolic compound PAA is known to have weak auxin activity and is widely distributed across the tree of life (Cook, 2019). Accordingly, it was omnipresent in our dataset. Endogenous IAA concentrations reported in chlorophyte and charophyte green algae, bryophytes, lycophytes and angiosperms are all similar in the way they vary across the entire nanomolar scale (**Table S4**); the values in our dataset were at the low-to-mid range of this spectrum.

Among cytokinins (CKs), we found N^6^-(Δ^2^-isopentenyl)-adenine (iP), *cis*-zeatin (cZ) and methylthio (MeS)-CKs to be the only types present in algae. Land plants additionally possessed *trans*-zeatin (tZ) and cZ/tZ-*O*-glucosides. Dihydrozeatin (DHZ) and CK-N-glucosides were absent altogether. Our results are consistent with the trend of dominance or exclusivity (as here) of tRNA-derived CKs in green algae and other lineages of living organisms (Stirk et al., 2003 & 2009 & 2013; Gajdosova et al., 2011; Zizkova et al., 2017; Seegobin et al., 2018; Aki et al., 2019 Pl Cell Physiol; Frebortova & Frebort, 2021). The non-tRNA CK types (tZ, DHZ) are generally detected much less consistently in green algae, varying (often within a single study) from none or trace to moderate amounts (Stirk et al., 2003 & 2009 & 2013; Ordog et al., 2004; Jiraskova et al., 2009; Hori et al., 2014; Zizkova et al., 2017). In vitro bryophytes appear to consistently contain tZ and no or little DHZ (von Schwartzenberg et al., 2007; Aki et al., 2019), though wild-collected material can sometimes deviate from this rule (Zaveska Drabkova et al., 2015). In angiosperms, CK can be deactivated by *O*- or *N*-glucosylation, which is reversible to varying degrees (Sakakibara et al., 2006; Hoyerova & Hosek, 2020; Pokorna et al., 2021). CK *N*-glucosides are only significantly accumulated in angiosperms (Ko et al., 2014; Zalabak et al., 2014; Prerostova et al., 2017; Vankova et al., 2017; Pokorna et al., 2021; Fuchs et al., 2022) and are almost universally absent in both algae (Stirk & van Staden, 2020) and bryophytes (Schwartzenberg et al., 2007 & 2016; Aki et al., 2019), in agreement with our results. Concerning CK *O*-glucosides, these have been reported at low concentrations in algae (Stirk et al., 2003 & 2009; Ordog et al., 2004; Zizkova et al., 2017), but are only significantly accumulated in land plants (compared to non-glucosylated CK forms), particularly the cZ-type O-glucosides in bryophytes (Zaveska Drabkova et al., 2015; Schwartzenberg et al., 2016; Aki et al., 2019) and several types in spermatophytes (Vankova et al., 2017; Zlobin et al., 2020; Fuchs et al., 2022). Our results fit this pattern only partially. We did not detect CK *O*-glucosides in algal material, but could confirm their presence in the three investigated land plants, although never in higher levels. The concentrations of CKs analyzed in this study were more or less comparable to those previously reported in green and other algae (Ordog et al., 2004; Stirk et al., 2003 and 2013; Jiraskova et al., 2009; Zizkova et al., 2017) and bryophytes (von Schwartzenberg et al., 2007; Lindner et al., 2014; Aki et al., 2019). Of note, seed plant vegetative tissues typically contain similar CK levels (Ko et al., 2014; Zalabak et al., 2014; Simura et al., 2016; Prerostova et al., 2017; Noah et al., 2021).

The jasmonate profile was particularly patchy in our dataset. No jasmonates were found in chlorophyte algae, while dinor-12-oxo-phytodienoic acid (dnOPDA) in *Marchantia polymorpha* (concentration similar to Monte et al. 2018) constituted the only jasmonate compound found in land plant samples. By contrast, some but not all tested charophytes contained jasmonic acid (JA), OPDA and dnOPDA in a partially overlapping fashion. Jasmonate-isoleucine (JA-Ile) was absent everywhere. Notably, dnOPDA (wherever found) far exceeded all other jasmonates in endogenous concentrations and was incredibly abundant in two strains of the charophyte species *Coleochaete scutata*. In all, it is counterintuitive to find multiple jasmonates in charophyte algae, but the literature is riddled with similar ostensible inconsistencies. The model bryophytes *Marchantia polymorpha* and *Physcomitrium patens* were found to contain OPDA and/or dnOPDA but not JA and JA-Ile (Stumpe et al., 2010; Ponce De León et al., 2012; Koeduka et al., 2015; Yamamoto et al., 2015), while other studies reported the presence of JA (Bandara et al., 2009; Oliver et al., 2009; Monte et al. 2018). Research on wild-collected bryophytes detected JA (Gachet et al., 2017) and even JA-Ile (Zaveska Drabkova et al., 2015). Similar controversies exist in lycophytes (Ogorodnikova et al. 2015; Gachet et al., 2017; Pratiwi et al., 2017; Monte et al., 2022; Wuyun et al., 2022). Among charophytes, no jasmonates were found in *Chara braunii* (Hackenberg & Pandey, 2014 and this study), in contrast to JA in wild-collected *Chara australis* (Beilby et al., 2015). Multiple studies on the same strain of *Klebsormidium nitens* NIES-2285 provide a more direct comparison: we only detected dnOPDA but no OPDA, whereas two other groups found both of these, and all three agree on JA-Ile absence (Koeduka et al., 2015; Monte et al., 2020). Additionally, in the same strain JA was both detected (Hori et al., 2014 and stationary cultures in this study) and not (Koeduka et al., 2015 and proliferative cultures in this study). To make matters more complicated, jasmonates were earlier reported in chlorophytes, red algae, cyanobacteria and even *Euglena* (Krupina & Dathe, 1991; Ueda et al., 1991a, 1991b; Fujii et al., 1997), prompting the hypothesis that jasmonate origin was associated with plastid endosymbiosis (Bouarab et al., 2004). Across land plants, jasmonate biosynthesis is highly responsive to stresses, with JA or OPDA endogenous levels increasing tens or even hundred(s) times after wounding or pathogen attack (Weber et al., 1997; Pan et al., 2008; Balcke et al., 2012; Scholz et al. 2012; Flokova et al., 2014; Pratiwi et al. 2017), whereas they may be present at low or even undetectable amounts under control conditions (Riedel et al., 2008; Radhika et al., 2012; Pratiwi et al. 2017; Resemann et al., 2023). This may account for the discrepancies in detecting endogenous jasmonates in unstressed land plants in this and other studies. Furthermore, *cis*- and dnOPDA have been implicated in a thermotolerance response that is most likely conserved between land plants and the charophyte *Klebsormidium nitens*, hence predating jasmonate signaling (Monte et al., 2020). In that regard, the variability in detection and endogenous levels of dnOPDA in charophytes in our dataset might simply reflect different temperature preferences among our algal strains.

Salicylic acid (SA) was detected in all tested species at varying concentrations ranging from low pmol to nmol/g FW, with no apparent difference between land plants and green algae. Previous studies found SA in chlorophytes and the charophytes *Chara braunii* and *Klebsormidium nitens*, generally at higher concentrations, although even in land plants the endogenous SA levels are known to vary (Hackenberg & Pandey, 2014; Hori et al., 2014; Jia et al., 2023 and references therein). Indeed, SA is apparently found ubiquitously in the green lineage (Jia et al., 2023), which is consistent with our results. Two main independent pathways for SA biosynthesis exist in plants, of which the ꞵ-oxidation-based is likely conserved in the green lineage (Jia et al., 2023). The direct SA precursor in this pathway is benzoic acid (BA), likewise detected in all strains examined in this study. However, a link between the two cannot be surmised based on endogenous detection alone. BA is a more general primary metabolite, important for the biosynthesis of other compounds and itself has multiple sources in plants (Widhalm & Dudareva, 2015). Therefore, BA-to-SA hydroxylation is the only step in the ꞵ-oxidation pathway specific to SA biosynthesis (Rieseberg et al., 2023). Although this step has been experimentally demonstrated in *Arabidopsis* (Wu et al., 2022), the putative hydroxylase has not been determined, as such a reaction could possibly be performed by a multitude of enzymes (Lefevere et al., 2020). Overall, the enzymatic machinery underlying SA biosynthesis is currently under investigation, even in established land plant models.

Although we included standards for certain gibberellins (GAs), brassinosteroids (BS) and strigolactones (SL), none were detected by our method in any of the tested organisms. This is not surprising, since these compounds are difficult to measure and typically necessitate dedicated approaches (Swaczynova et al., 2007; Urbanova et al., 2013; Li et al., 2016; Deng et al., 2016; Yoneyama et al., 2018; Walker et al., 2019). That said, some compounds included in our analyte screen were reported in organisms equivalent or closely related to those in our sample set. GA4 and GA24 were found in *Selaginella moellendorffii,* which contains canonical GAs biosynthesis and response (Hirano et al., 2007; Yasumura et al., 2007; Hernandez-Garcia et al., 2021). GA3 presence was suggested in the antheridia of two *Chara* species (Kazmierczak & Stepinski, 2005). Multiple GAs and the BS brassinolide and carlactone were detected in unicellular chlorophytes (Stirk et al., 2013 Physiol Biochem 70), with *Chlorella vulgaris* even reported to contain 7 BS (Bajguz, 2009). Out of the three SL in our analyte list, all were previously reported to be excreted by *Physcomitrium patens* protonema (Proust et al., 2011) and two were produced in *Marchantia polymorpha* thallus (Delaux et al., 2012).

### Phytohormone levels in culture and control media

The potential relevance of extracellular release of phytohormones has been debated in organisms that lack complex tissues or differentiated cell types to plausibly utilize phytohormone gradients (Hartung et al., 2010; Bennett et al., 2015; Vosolsobe et al., 2020). Strigolactones were implicated in quorum sensing in *Physcomitrium patens* protonema (Proust et al., 2011). To investigate the extent of extracellular phytohormone release by the investigated organisms, we mirrored our biomass screen by the same analysis in the corresponding culture media.

Auxins, CKs and phenolics (SA, PAA, BA) were present in the culture media (**Figure 2A,B**). CKs were nearly universally less abundant in media than in biomass, whereas this was variable for the other phytohormone types (**Figure 2A**). Comparing the values between culture and blank media revealed that the blanks universally lacked IAA and typically did not contain CKs, whereas the relationship was again variable in regard to phenolics (**Figure 2B**). ABA and jasmonates were absent in blanks and could not be found in culture media either, except for some charophyte strains in stationary phase, which also contained these compounds in biomass (**Figure 1, Figure 2A,B**). IAA excretion in *Physcomitrium patens* protonema was previously linked to the action of the PIN-FORMED family auxin efflux proteins (Viaene et al., 2014). *Klebsormidium* spp. was shown to contain a functionally conserved homolog and likewise excretes IAA (Skokan et al., 2019 and this study). However, *Coleochaete scutata* also excreted IAA (**Figure 2A,B**), although no PIN-FORMED homologs were found in this lineage (Vosolsobe et al., 2020), underscoring this protein family is not necessary to exude IAA. The chlorophyte algae (including *Chlorella* spp.) were previously found to excrete IAA and ABA (Marsalek et al., 1992; Mazur et al., 2001; Prieto et al., 2011; Khasin et al., 2018; Pichler et al., 2020). We did not detect these compounds in the culture media of the 4 chlorophytes studied, except for IAA in the stationary phase *Chlorella sorokiniana* (**Figure 2A,B**). CK excretion is apparently widespread in the tree of life (Stirk & van Staden, 2010) and accordingly all CK types produced in biomass were also found in the corresponding culture media in this study, interestingly including tZ-types wherever these were found.

**Figure 2.**
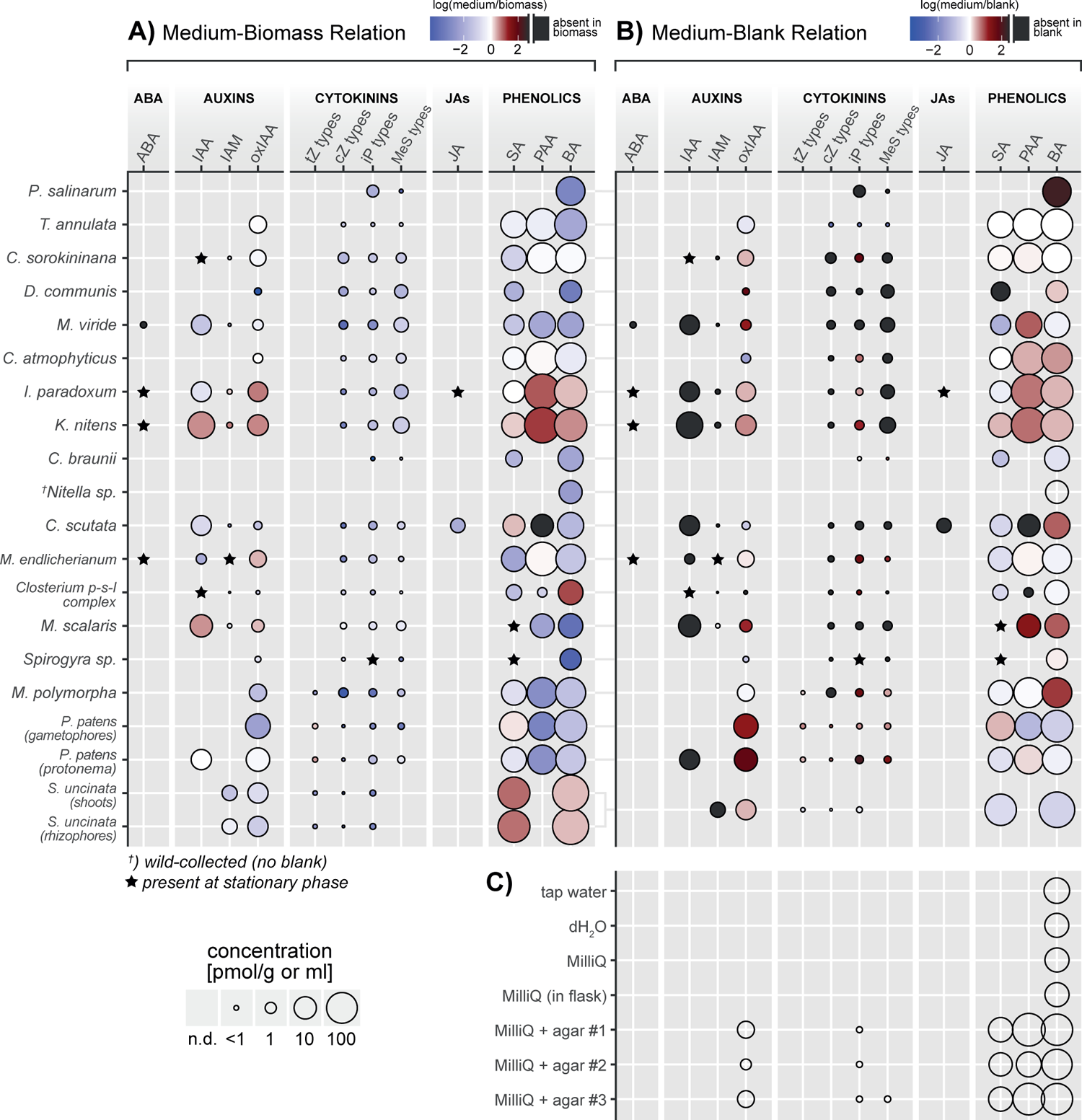
Detection of phytohormone compounds in culture media and control samples of water and agar. **A)** Ratio between concentrations detected in culture medium and the corresponding biomass, in logarithmic scale. Color code: Blue–compound prevalent in biomass. Red–compound prevalent in medium. Black–compound absent in biomass. **B)** Ratio between concentrations detected in culture medium and blank medium (containing no biological material), in logarithmic scale. Color code: Blue–compound prevalent in blank. Red–compound prevalent in medium. Black–compound absent in blank. **C)** Compounds detected in different purity grades of water and three brands of agar (1.5% w/v, incorporated in MilliQ grade water). The circle size in (A,B,C) denotes concentration in media (pmol per gram or ml). No circle–compound not detected (n.d.). Compound abbreviations are listed in the legend to Figure 1.

To address possible environmental contaminants, we additionally analyzed different purity grades of water and three brands of agar (**Figure 2C**). BA was the only compound detected in water. However, due to its chemical nature, BA was the only analyte lacking a qualifier ion, so the signal might not be specific to BA. By contrast, the samples of agar (incorporated in MilliQ grade water) contained oxIAA, iP and considerable amounts of the phenolics SA and PAA (in addition to BA). This is likely due to the biological source of agar. Agarophyte red algae are known to produce IAA, CKs and SA (Yokoya et al., 2010; Farvin & Jacobsen, 2013; Hou et al., 2018). The majority of IAA could putatively become oxidized during the process of agar production. The concentrations of SA measured in agar (10^1^ nM) do not appear insignificant compared to those found in biomass and culture media, but are still far below the effective doses of applied SA in land plants (Cao et al., 1994; Beyer et al., 2021).

The issue of culture axenicity in phytohormone measurements and treatments in algae has long been debated (Evans & Trewavas, 1991; Cook, 2019). Three charophyte species in this analysis were available for comparison as both axenic and contaminated with bacteria. However, the phytohormone profiles between them were qualitatively nearly identical (**Figure S3A**). As for the culture medium, some differences were observed, but these were not uniform (**Figure S3B**). For instance, the medium of the contaminated *Chlorokybus atmophyticus* contained IAA, whereas that of the axenic strain did not. However, *Mesostigma viride* and *Coleochaete scutata* showed the opposite pattern. Apparently, the effects of bacteria and algae co-cultivation might be strain-specific to either or both organisms. Although the effects of contamination were not as significant as might have been expected, it is still highly advisable to avoid contamination whenever possible.

### Inconsistencies in plant hormonomics

The efforts to determine endogenous phytohormone concentrations have spanned decades. The general picture is complicated even in angiosperms, where the majority of phytohormone research has been performed. Different methods, plant species, organs, developmental stages and conditions in culture and experiments have understandably led to varying results. Applying two methods to the same material within one study could bring both very similar (Dobrev et al., 2005; Kojima et al., 2009) and different (Jayaswal & Johri, 1985; Balcke et al., 2012) results in the levels of analytes measured. While different organs of the same plant understandably have different phytohormone levels (e.g. Muller et al., 2002; Veach et al., 2003), the reported control levels even in the same organ (and same species) can vary by an order of magnitude between studies. Examples include ABA (Pan et al., 2008; Flokova et al., 2014; Prerostova et al., 2021; vs. Farrow & Emery, 2012; Vankova et al., 2017; Muller et al., 2002), IAA (Flokova et al., 2014; Prerostova et al., 2021; Vankova et al., 2017 vs. Pan et al., 2008; Farrow & Emery 2012; Izumi et al., 2009) and JA (Pan et al., 2008; Flokova et al., 2016; Prerostova et al., 2017 & 2021 vs. Muller et al., 2002; Schmelz et al., 2003 & 2004; Vankova et al., 2017) in Arabidopsis leaves, JA-Ile in Arabidopsis roots (Prerostova et al., 2017; Simura et al., 2018 vs. Vankova et al., 2017; Prerostova et al., 2017) or cZ in maize shoots (Zalabak et al., 2014 vs. Veach et al., 2003). High variation can be observed in the same tissue and species at slightly different developmental stages. For instance, seedlings aged 1-3 days apart can show drastic differences in the concentrations of IAA metabolites (Tam et al., 2000; Noah et al., 2021) or tZ (Skalak et ak., 2019), yet the “seedling” material in different studies often differs in days after germination, sometimes with little or no additional information provided. The same issues occur in the comparatively less studied bryophytes and green algae. Quite different control IAA concentrations have been reported in protonema of the moss *Funaria hygrometrica* (Jayaswal & Johri, 1985 vs. Atzorn et al, 1990; Bhatla, 1992). The species/strains within the chlorophyte genus *Chlorella* differ significantly in the measured endogenous concentrations of ABA (Marsalek et al., 1992; Stirk et al., 2014), IAA (Stirk et al., 2013 & 2014) and CKs (Ordog et al., 2004; Jiraskova et al., 2009; Stirk et al., 2013 & 2014). The same is true for other species/strains of chlorophyte microalgae (Ordog et al., 2004; Stirk et al., 2013; Zizkova et al., 2017; Gonzales Cruz et al., 2023). In the charophyte genus *Chara*, three studies differ substantially in endogenous ABA and IAA concentrations (Hackenberg & Pandey, 2014; Tietz et al., 1989; Beilby et al., 2015). In all, we can see that differences in methods, experimental material and culture/experimental conditions inevitably lead to variance in measured endogenous phytohormone levels, and this problem is compounded when limited data are available, such as in charophytes or bryophytes.

It seems more appropriate to employ qualitative comparisons (presence vs. absence of particular compounds) where evolutionary distances between lineages are considerable. Even this approach is not without drawbacks, especially when the measured concentrations approach the detection limit. Hence ABA was both found and not found (Rathnayake et al., 2018 vs. Xiao et al., 2018 and this study) in ‘unstressed’ *Physcomitrium patens* protonema and the chlorophyte genus *Draparnaldia* (Tietz et al., 1989 vs. Tietz et al., 1986). JA was similarly detected (Beilby et al., 2015) and not detected (Hackenberg & Pandey, 2014) in the charophyte genus *Chara*. Three previous studies on the charophyte genus *Klebsormidium* differ in both detection and endogenous levels of different CKs, while simultaneously closely agreeing on the concentration of IAA (Hori et al., 2014; Stirk et al., 2013; Zizkova et al., 2017).

To complicate the issue further, green algae and bryophytes typically do not contain certain phytohormone metabolites, but can readily produce them when supplied with an appropriate precursor. While *Chlorella minutissima* (Stirk et al., 2014) and all green algae in this study lacked the ABA catabolites PA & DPA, the chlorophyte genus *Dunaliella* contained fairly high endogenous ABA concentrations (Tietz et al., 1989) and was able to convert exogenously applied ABA to PA and DPA (Cowan & Rose, 1991), although this may be related to its rich carotenoid metabolism (Xu & Harvey, 2019). IAA conjugates in the model bryophytes *Marchantia polymorpha* and *Physcomitrium patens* may be below the detection limit under normal conditions, but are significantly accumulated after exogenous IAA treatment (Kaneko et al., 2020). Similar metabolization of applied IAA was observed in the charophyte genus *Nitella* (Sztein et al., 2000) and chlorophyte microalgae (Zizkova et al., 2017). The cytokinin tZ was not found in two strains of the charophyte genus *Klebsormidium* (Stirk et al., 2013; Hori et al., 2014 and this study), while a third strain contained a small amount of tZ and could catabolize exogenously applied tZ (Zizkova et al., 2017). We are hence reminded to appreciate the unexplored and likely significant and divergent metabolic potential in green algae (Rieseberg et al., 2023) and to keep in mind the potentially situation-dependent nature of phytohormone compound biosynthesis and metabolism. The community should strive to observe as much uniformity as possible in experimental conditions and methodology, which must be described in detail, including but not limited to illumination spectra and ratios of fresh to dry weight, in case the latter is used.

### Evolutionary implications of phytohormone profiling

Before drawing conclusions from our data and the available literature, we briefly mention that conciliating phylogenomic and metabolomic evidence can be complicated. One example is the Gretchen Hagen 3 (GH3) enzymes that produce IAA- and JA-amino acid conjugates in land plants (Casanova-Saez et al., 2021). The GH3 family is absent in green algae (Bowman et al., 2021), yet multiple species were reported to convert applied IAA to IAA-amides (Sztein et al., 2000; Zizkova et al., 2017). On the other hand, the charophyte strain *Klebsormidium nitens* NIES-2285 acquired GH3 by horizontal gene transfer (Bowman et al., 2021), yet did not contain detectable endogenous IAA-amides in this study. A homolog of the GH3 member JAR1 was identified in the transcriptome of *K. crenulatum* (Holzinger et al., 2014), although studies of *K. nitens* NIES-2285 revealed no endogenous JA-Ile (Hori et al., 2014; Koeduka et al., 2015 and this study). The UGTs recognized for IAA conjugation to glucose are restricted to seed plants (Wilson & Tian, 2019), yet we measured IAA-GE in *Marchantia polymorpha* and *Selaginella uncinata*. The cytokinin oxidase/dehydrogenase (CKX) enzymes responsible for irreversible CK degradation are absent in green algae (Dabravolski & Isayenkov, 2021). Accordingly, several chlorophytes and the charophyte *Klebsormidium flaccidum* were shown to lack detectable CKX activity, yet they could still efficiently cleave applied tZ into adenosine and adenine (Zizkova et al., 2017). These facts point out the limitations of evolutionary inferences based primarily on comparative genomics, particularly when land plants (or angiosperms) comprise the bulk of available knowledge.

While phytohormone profiling in green algae has mostly focused on unicellular chlorophytes (Han et al., 2018; Stirk & van Staden, 2020; Wang et al., 2022), it is peculiar that these apparently produce almost all known phytohormones, and often in richer spectra than charophytes or even bryophytes (Hirano et al., 2007; Stirk et al., 2013a,b; Hori et al., 2014; Zaveska Drabkova et al., 2015; Zizkova et al., 2017; Aki et al., 2019). Given the aforementioned variability in results between studies, one wonders how all of these reports would compare if all organisms were subjected to a single analysis. Considering the vast evolutionary rift between land plants and organisms as distant as chlorophytes or more (Bowles et al., 2022), a kind of conserved phytohormone response seems to be one of the less likely imaginable explanations for observations such as evidence of endogenous phytohormones, a culture growth or metabolic response to their external application, or the presence of some phytohormone-related gene orthologs.

Minding everything discussed so far, we favor the view that most of the compounds recognized as phytohormones only became the eponymous developmental regulators in the ancestral land plant. Auxin and ABA are perhaps the clearest examples. Their developmental roles in bryophytes are well recognized, with significant phenotypic and physiological responses reported for treatments at lower micromolar doses (Nagao et al., 2005; Yasumura et al., 2007; Flores-Sandoval et al., 2015; Lavy et al., 2016; Eklund et al., 2018; Mutte et al., 2018; Thelander et al., 2018; Jahan et al., 2019; Suzuki et al., 2022). This is not observed In charophytes or much higher doses are required to elicit a notable response (Klambdt et al., 1992; Sederias & Colman, 2007; Nagao et al., 2008; Ohtaka et al., 2017). The canonical auxin and ABA receptors are present and functional in bryophytes (Mutte et al., 2018; Jahan et al., 2019; Sun et al., 2020; Suzuki et al., 2022), whereas in charophytes they are either absent (auxin) (Bowman, 2021) or do not bind the hormonal ligand (ABA) (Sun et al., 2019). Accordingly, we found that land plants are clearly distinguished from charophytes by the consistent detection of ABA and the presence of auxin conjugates, but also by the presence of CK conjugates and tZ. In fact, both auxin and CK profiles of green algae resemble those of non-green organisms (Morrison et al., 2015; Hendrikx & Schnabl, 2019; Chen et al., 2022; **Table S7**) including animal tissues (Seegobin et al., 2018; Aoki et al., 2019; Liu et al., 2022), where these reflect tryptophan/indole metabolism and the deeply conserved tRNA modifications, respectively (Dabravolski et al., 2020; Gibb et al., 2020). Indeed, our results suggest that CK response is also a land plant invention, but this argument is less compelling. Not all studies conclude that the CK profiles of green algae and bryophytes differ that much (e.g. Stirk et al., 2013; von Schwartzenberg et al., 2016), although CKs clearly regulate development in the latter (Rashotte et al., 2021). The two-component CK signaling is conserved between land plants and charophytes, but the orthologs of the charophyte lineage Zygnematophyceae are not significantly transcriptionally responsive to CK treatments (de Vries et al., 2020). Further research is needed to identify clear hallmarks of CK response. In the case of jasmonates and SA, these compounds were found in both charophytes and land plants, although the canonical receptors only emerged in the latter (Schluttenhofer, 2020; Jia et al., 2023). It is conceivable that some phytohormones had other physiological role(s) before they were integrated into the recognized canonical signaling pathways and that this might have been retained and modified in some form by certain extant charophytes. A role of the jasmonate dnOPDA in thermotolerance appears to be one such case (Monte et al., 2020). It likewise seems plausible that certain conditions lead to an increased production of compounds that would eventually come to regulate the physiological response to such conditions as phytohormones. Such a scenario was proposed for ABA resulting from increased carotenoid oxidation under stress (Rieseberg et al., 2023), concurrent with observations in green algae that intracellular ABA content may increase under stress, but exogenous ABA treatment does not affect stress responses (Takezawa et al., 2011; Sun et al., 2020).

The extant diversity in the streptophyte clade is a testament to the many different evolutionary paths taken by its individual lineages. We hope that this study could serve as one of the many steps that need be taken to unravel the many tumultuous changes that occurred during the transition to land in the ancient organisms now lost to us.

## MATERIALS & METHODS

### Chemicals

**Table S5** lists all chemicals used in this study.

### Algal & plant strains and cultivation

Algal strains were obtained from the following institutions: Microbial culture collection, National institute for environmental studies, Tsukuba, Japan (NIES); Culture collection of algae, Goettingen University, Germany (SAG); Central collection of algal cultures, Duisburg University, Essen, Germany (CCAC); Culture collection of algae, University of Texas at Austin, USA (UTEX); Culture collection of algae, Department of Botany, Charles University, Prague, Czechia (CAUP). *Selaginella uncinata* (“Comenius”, original strain) was obtained from the Botanical garden at the Comenius University, Bratislava, Slovakia, and established as a sterile culture. The wild-collected samples of *Nitella* sp. and the surrounding water (flash-frozen in liquid N_2_ *in situ*) were obtained from a freshwater spring pond under sandstone rocks (GPS: 50.6491178N, 14.5136108E) in April 2021; the surrounding water was notably pure and the algae not obviously covered by epiphytic microflora.

See **Table S1** for a complete list of strains with source identifiers and the culture media used. Murashige-Skoog medium (Duchefa M0221), Gamborg medium (Duchefa G0210) and Bold’s Basal Medium (BBM; Merck B5285) were purchased commercially. BCD medium supplemented with diammonium tartrate (BCDAT) was prepared according to Cove et al., 2009. C medium (Ichimura, 1971), Pro medium (Starr, 1964; Ichimura & Itoh, 1977) and a modified version of the SWCN-4 medium for Charophyceae were prepared per instructions from the NIES collection (https://mcc.nies.go.jp/02medium-e.html). Modified SWCN-4 as follows: garden soil was mixed with river sand (1:4), dampened with dH_2_O, autoclaved 3x, laid into sterile test tubes (3 cm height) and supplemented with 40 ml sterile dH_2_O. BBM, C and Pro media were supplemented with customized vitamin doses: B_12_ 10 mg/l medium, B_7_ 2 mg/l, B_1_ 10 mg/l. Individual vitamin stocks (1000x) were dissolved in dH_2_O, sterilized by filtration (Millex SLGS033SS), stored at −20°C, and added into cooled-down but not yet solid agar-supplemented medium, or upon algal inoculation (liquid medium). All media were sugar-free.

Land plants were cultured on solid media in plates sealed with surgical tape (Micropore 1530-0). *Marchantia polymorpha* were inoculated from individual gemmae. *Physcomitrium patens* protonema were homogenized weekly using Ika T25 Digital Ultra Turrax with 8G disperser tool (IKA-Werke, Germany) and spread on plates containing medium overlaid with cellophane foil. When necessary, cultures were left to grow without homogenization to allow gametophore emergence. *Selaginella uncincata* (chosen for its superb growth *in vitro*) was inoculated from apical branch cuttings. 2-3 apical explants of *Chara braunii* (each with at least two nodes) were inoculated per one test tube containing soil and culture medium. Other algae, if not grown for analysis, were inoculated into fresh medium from a fraction of the original culture (2-10% biomass) in 6-8 week intervals or longer, depending on the growth character of each individual strain.

Although we have generally avoided this measure in cultures intended for phytohormone analysis, certain algal strains (**Table S1**) require supplementation of the culture media with soil extract for continuous proper growth. Soil extract was prepared by mixing soil with dH_2_O (1:3 volume), autoclaving, passing through filter paper overnight and autoclaving again. The soil (without obvious leaf litter) was collected in a submontane forest of European beech (*Fagus sylvatica*; GPS: 50.8653578N, 15.1046144E) at the end of winter.

All living material was cultured at 23°C, 16:8 hours light:dark regime. The cultures were illuminated by mixed-spectrum fluorescent light tubes, using Osram Fluora TLD 36W (Osram Licht AG, Germany) and Philips Master TLD Super 36W (Koninklijke Philips N.V., The Netherlands); see **Table S6** for details on illumination spectra and intensity.

### Sample preparation

Algal biomass was sampled during the proliferative phase of culture growth (1.5-3 weeks after inoculation, depending on individual strains). Streptophyte algae were additionally sampled during the stationary phase of growth (4-6 weeks). *Marchantia* thalli were sampled on day 19. *Physcomitrium* protonema and gametophytes/thalli/plants were sampled after 1 and 5 weeks of growth, respectively. *Selaginella* was sampled 5 weeks post inoculation.

Samples were harvested as follows: liquid-cultured algae were collected by pipetting (cut-off tips) and filtration through a nylon mesh filter (20 μm, Merck Millipore NY2004700) using underpressure. Certain strains (**Table S1**) were collected by centrifugation (1000 x g) and supernatant decantation. Algae and moss protonema growing on solid media were scraped with a spatula (and filtered to remove excessive moisture, if necessary). *Chara braunii* and *Nitella* sp. thalli (without rhizoids) were washed with dH_2_O and excessive moisture removed with a cotton pad. Bryophyte gametophores and *Selaginella* apical cuttings were simply transferred into collection tubes. Biomass samples (10-50 mg fresh weight each) were transferred into 2-ml thick-walled microcentrifuge tubes (SSIbio 2340-00, Scientific Specialities, USA) with screwable lids (SSIbio 2002-00), flash-frozen in liquid nitrogen and stored at −80°C. Liquid culture media were sampled by pipetting, centrifugation (1000g) and decanting as 100 µl samples. Samples of solid culture media were scraped from inside the solid agar (avoiding any residual biomass) with a cut pipette tip and sampled as ca. 50 mg each. Blank media were sampled similarly. The media samples were likewise flash-frozen in liquid nitrogen and stored at −80°C.

### Phytohormone analysis

Samples (biomass and solidified media ca. 10-50 mg FW, liquid media 100ul) were extracted with 100 µl 1M formic acid solution. The following isotope-labelled standards were added at 1 pmol per sample: ^13^C_6_-IAA (Cambridge Isotope Laboratories, Tewksbury, MA, USA); ^2^H_4_-SA (Sigma-Aldrich, St. Louis, MO, USA); ^2^H_3_-PA, ^2^H_3_-DPA (NRC-PBI); ^2^H_6_-ABA, ^2^H_5_-JA, ^2^H_5_-tZ, ^2^H_5_-tZR, ^2^H_5_-tZRMP, ^2^H_5_-tZ7G, ^2^H_5_-tZ9G, ^2^H_5_-tZOG, ^2^H_5_-tZROG, ^15^N_4_-cZ, ^2^H_3_-DZ, ^2^H_3_-DZR, ^2^H_3_-DZ9G, ^2^H_3_-DZRMP, ^2^H_7_-DZOG, ^2^H_6_-iP, ^2^H_6_-iPR, ^2^H_6_-iP7G, ^2^H_6_-iP9G, ^2^H_6_-iPRMP ^2^H_2_-GA19, (^2^H_5_)(^15^N_1_)-IAA-Asp and (^2^H_5_)(^15^N_1_)-IAA-Glu (Olchemim, Olomouc, Czech Republic). The extracts were centrifuged at 30,000×g at 4°C. The supernatants were applied to SPE Oasis HLB 96-well column plates (10 mg/well; Waters, Milford, MA, USA) conditioned with 100 µL acetonitrile and 100 µl 1M formic acid using Pressure+ 96 manifold (Biotage, Uppsala, Sweden). After washing the wells three times with 100 µl water, the samples were eluted with 100 µl 50% acetonitrile in water. The eluates were separated on Kinetex EVO C18 HPLC column (2.6 µm, 150 × 2.1 mm, Phenomenex, Torrance, CA, USA). Mobile phases consisted of A—5 mM ammonium acetate and 2 µM medronic acid in water and B—95:5 acetonitrile:water (v/v). The following gradient was applied: 5% B in 0 min, 5–7% B (0.1–5 min), 10– 35% B (5.1–12 min) and 35–100% B (12–13 min), followed by a 1 min hold at 100% B (13–14 min) and return to 5% B. Hormone analysis was performed with a LC/MS system consisting of UHPLC 1290 Infinity II (Agilent, Santa Clara, CA, USA) coupled to 6495 Triple Quadrupole Mass Spectrometer (Agilent, Santa Clara, CA, USA), operating in MRM mode, with quantification by the isotope dilution method. Data acquisition and processing was performed with Mass Hunter software B.08 (Agilent, Santa Clara, CA, USA).

## ACKNOWLEDGMENTS

We acknowledge Pavel Škaloud (Department of Botany, Charles University, Prague, Czechia) for providing the CAUP algal cultures and valuable advice; Norbert Zlámal (Botanical Garden, Comenius University, Bratislava, Slovakia) for identifying and providing *Selaginella uncinata*; Marie Korecká (Institute of Experimental Botany, Czech Academy of Sciences, Prague, Czechia) for assistance with experimental work. This study was supported by Czech Science Foundation project 20-13587S.

## COMPETING INTERESTS

The authors declare no competing interests.

## Notes

### Competing Interest Statement

The authors have declared no competing interest.

