## Supplementary material for "Phytohormone profiling in an evolutionary framework": Schmidt-Skokan-et-al-2023_supp-figures.pdf


#### AUTHORS & AFFILIATIONS

Vojtěch Schmidt[1,2†], Roman Skokan[1†*], Katarina Kurtović[2], Stanislav Vosolsobě[2], Roberta Filepová[1], Samuel Haluška[1], Petre Dobrev[1], Václav Motyka[1], Jan Petrášek[1,2*]

1) Institute of Experimental Botany of the Czech Academy of Sciences, Rozvojová 263, Prague, Czechia
2) Department of Experimental Plant Biology, Charles University, Viničná 5, Prague, Czechia

†) co-first authors
*) corresponding authors:
Roman Skokan,
Jan Petrášek,


### Supplementary Material

##### Supplementary Figures (this file)

Figure S1. Endogenous phytohormone compounds detected in the biomass of green algae and land plants.
Figure S2. Endogenous phytohormone compounds detected in biomass and culture media of selected green algae, stationary vs. proliferative cultures.
Figure S3. Phytohormone detection in axenic and contaminated strains.

##### Supplementary Tables (separate file)

Table S1. Strain list.
Table S2. Analyte list.
Table S3. Raw Data.
Table S4. Phytohormone levels in literature.
Table S5. Chemical list.
Table S6. Illumination.
Table S7. Phytohormone production by digestion via gut bacteria.

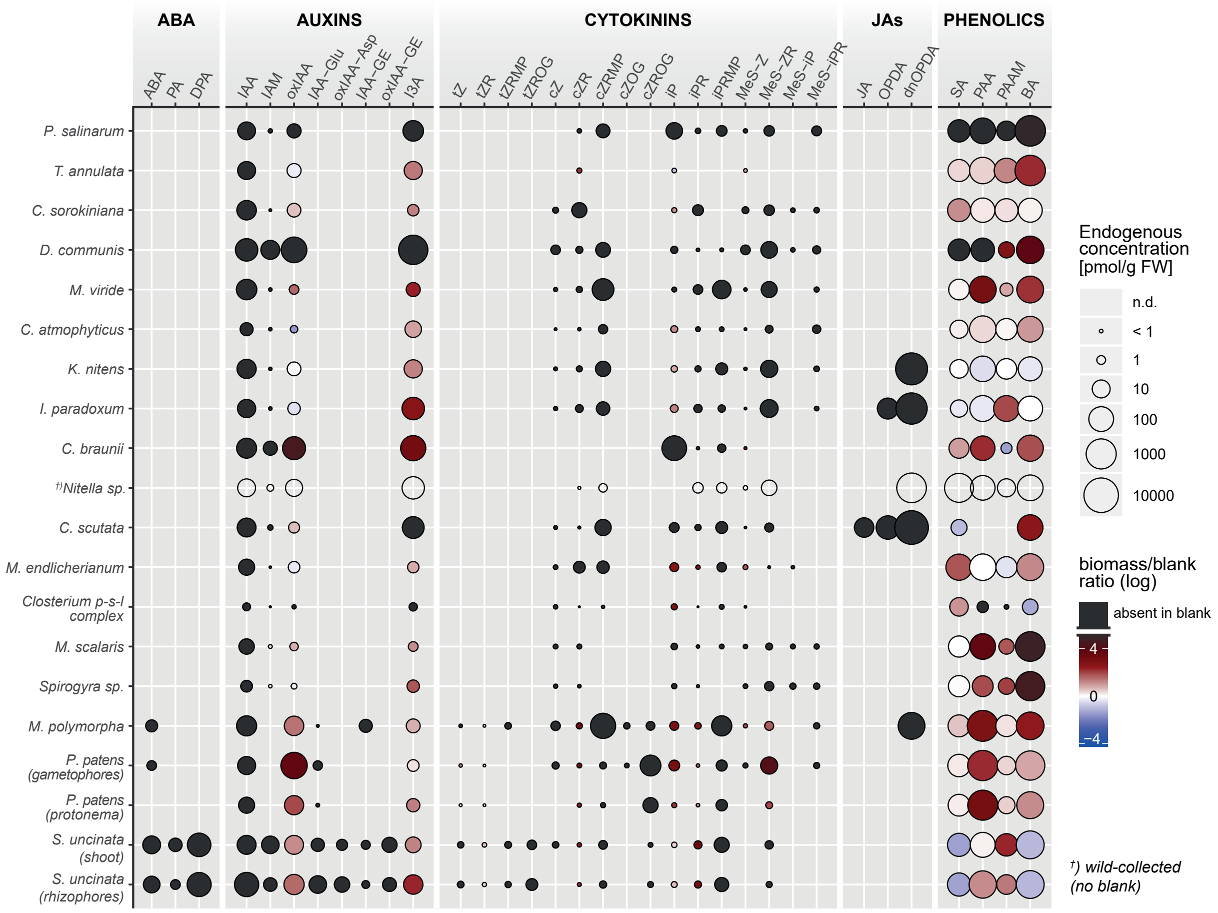


**Figure S1. Endogenous phytohormone compounds detected in the biomass of green algae and land plants.** Corresponds to Figure 1, but all analytes detected are listed. Circle size denotes concentration (pmol per gram fresh weight). No circle–compound not detected (n.d.). Color code denotes the ratio between the values measured in biomass and blank medium (containing no biological material), expressed in logarithmic scale. Blue–compound(s) prevalent in blank. Red–compound(s) prevalent in biomass. Black–compound(s) absent in blank. Abbreviations: ABA (abscisic acid), PA (phaseic acid), DPA (dihydrophaseic acid), IAA (indole-3-acetic acid), IAM (indole-3-acetamide), oxIAA (2-oxo-IAA), IAA-Glu (IAA-glutamate), IAA-GE (IAA-glucose ester), tZ (*trans*-zeatin), cZ (*cis*-zeatin), Z(R)OGs (zeatin (riboside)-O-glucosides; both *cis*- and *trans*-isomers), iP (N6-(Δ2-isopentenyl)-adenine), MeS (methylthio), JA (jasmonic acid), OPDA (12-oxo-phytodienoic acid), dnOPDA (dinor-OPDA), SA (salicylic acid), PAA (phenylacetic acid), BA (benzoic acid).

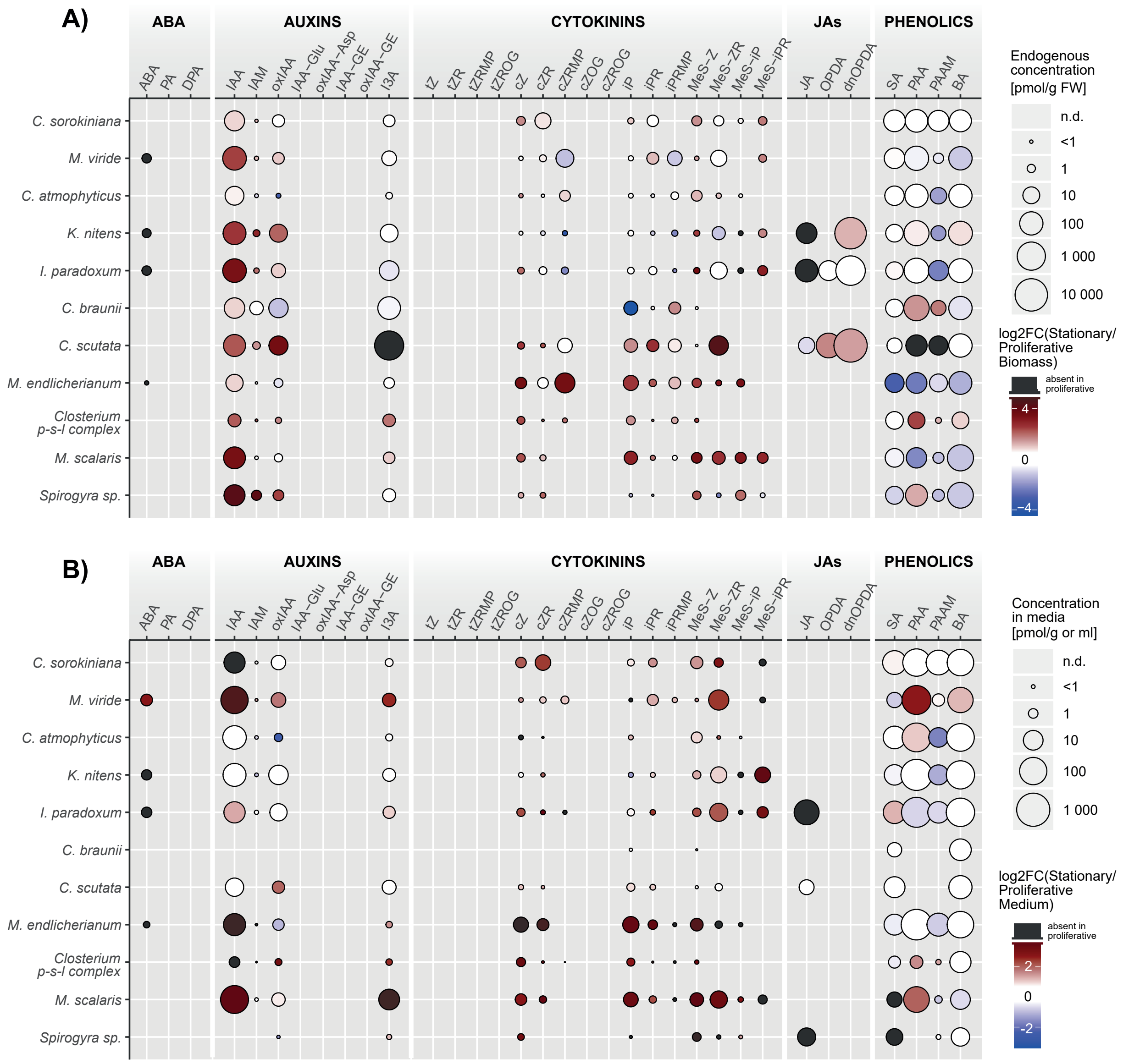


**Figure S2. Endogenous phytohormone compounds detected in biomass and culture media of selected green algae, stationary vs. proliferative cultures. A)** Ratio between biomass in stationary and proliferative growth phases, logarithmic scale. Color code: Blue–prevalent in proliferative phase. Red–prevalent in stationary phase. Black–absent in proliferative phase. Circle size denotes concentration in stationary cultures (pmol per gram fresh weight). **B)** Same as A), but in culture media. Circle size denotes concentration in stationary culture media (pmol per gram or ml). No circle in (A,B): compound not detected (n.d.).

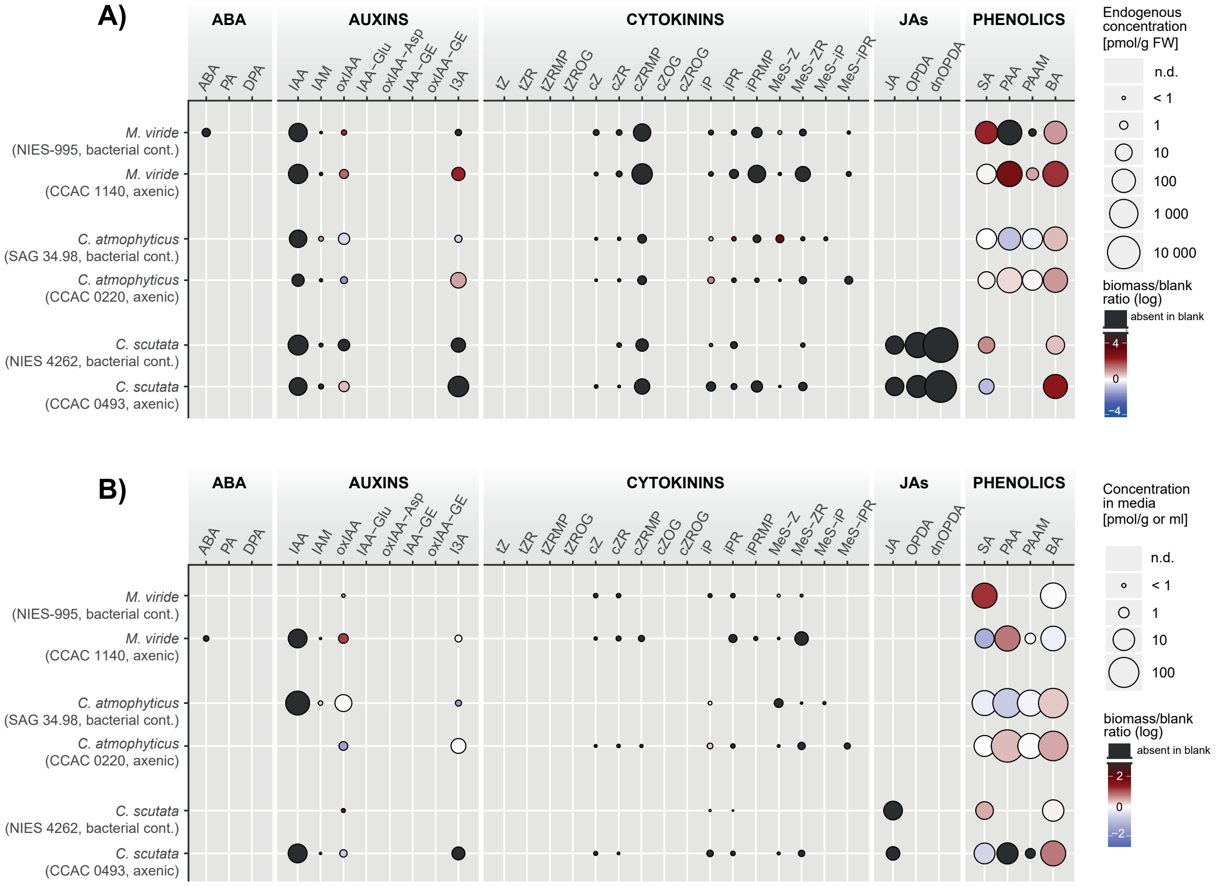


**Figure S3. Phytohormone detection in axenic and contaminated strains. A)** Ratio between the values measured in biomass and blank medium (containing no biological material), expressed in logarithmic scale. Color code: Blue–prevalent in blank. Red–prevalent in biomass. Black–absent in blank. Circle size denotes concentration in biomass (pmol per gram fresh weight). **B)** Ratio between the values measured in culture media and corresponding blank media. Color code: Blue–prevalent in blank. Red–prevalent in culture medium. Black–absent in blank. Circle size denotes concentration in culture media (pmol per gram or ml). No circle in (A,B): compound not detected (n.d.).
